# MEEGNet: An open source python library for the application of convolutional neural networks to MEG

**DOI:** 10.1101/2025.03.20.644276

**Authors:** Dehgan Arthur, Pascarella Annalisa, Yann Harel, Rish Irina, Jerbi Karim

## Abstract

Artificial Neural Networks (ANNs) are rapidly gaining traction in neuroscience, proving invaluable for decoding and modeling brain signals from techniques such as electroencephalography (EEG) and functional magnetic resonance imaging (fMRI). Although these networks are beginning to find applications in magnetoencephalography (MEG), their use in this domain is still in the early stages. Here, we introduces MEEGNet, a novel Python library paired with an intuitive convolutional neural network (CNN) architecture designed primarily for MEG data, yet adaptable to EEG signals. The MEEGNet model was trained and cross-validated using MEG data from 643 participants across four classification tasks, including auditory and visual stimulus classification and age prediction. Our model achieves competitive performance across all tasks, with a notable balance of accuracy and efficiency—for instance, reaching 92.70% test accuracy in an auditory vs. visual classification task while maintaining shorter training times than other architectures. The MEEGNet pipeline also integrates latent space visualization tools, adapted for MEG and EEG data. These include saliency maps and Grad-CAM methods, which enhance the interpretability of ANN-based classification and help address the black-box critique of such models. Importantly, the MEEGNet library is designed for extensibility, allowing the neuroscience and machine learning communities to add functionalities and ANN models. By prioritizing usability, transparency, and interpretability, MEEGNet empowers MEG- and EEG-based research with a user-friendly and modular framework.

## Introduction

Artificial Neural Networks (ANNs), including Convolutional Neural Networks (CNNs), have become pivotal tools across many research domains, transforming tasks like image recognition, natural language processing, and predictive modeling [1]. Their capacity to uncover complex patterns has recently gained traction in neuroscience, particularly for decoding brain signals (For a review see [2]).

There is a clear surge in interest towards leveraging ANNs for the analysis of brain data, yet their adoption by the MEG and EEG community is still relatively slow. Tools such as MNEFlow [3] and Braindecode [4] exemplify the burgeoning suite of resources catering to this enthusiasm. However, the complexity and steep learning curve associated with artificial neural networks (ANNs), combined with a shortage of well-documented, specialized open source frameworks, frequently compel researchers to opt for simpler alternatives like scikit-learn machine learning models. Moreover, the challenge of interpreting ANN models, which is critical for deriving scientific and clinical insights, is still an underexplored area in magnetoencephalography (MEG) studies.

In an attempt to bridge these gaps, this paper introduced MEEGNet, an open-source Python library designed to simplify ANN application to MEG and EEG decoding tasks, with an emphasis on usability, interpretability, and computational efficiency. Building on the growing use of ANNs in M/EEG research, MEEGNet provides streamlined data processing, training, and evaluation tools, alongside pre-trained models and data augmentation features. It includes established ANN architectures adapted for M/EEG data, such as MLP [5], AlexNet [6], VGG-16 [7], EEGNet [8] and our own custom-designed CNN architecture tailored for MEG data. A key innovation is the integration of latent space visualization tools, like Guided Backpropagation Saliency maps, tailored to M/EEG signals. These built-in visualization tools were developed to allow for a deeper understanding of network activations and their correspondence with brain dynamics. By combining all these capabilities, MEEGNet lowers entry barriers, enhances transparency, and fosters reproducible ANN-driven MEG analysis.

## Design and Implementation

Below, we outline the core components of the MEEGNet library, focusing on its primary classes and functions. For a detailed exploration of these elements, see the API section of the online documentation or review the included tutorials.

### Data Loading

The MEEGNet library provides a robust data-loading functionality through a hierarchical class structure featuring two primary classes: `EpochedDataset` and `ContinuousDataset`. The `EpochedDataset` class is optimized for event-based data, where recordings are segmented around specific events, such as stimulus presentations. In contrast, the `ContinuousDataset` class handles continuous data without event markers, such as resting-state recordings. While these classes are tailored for specific data types, users can flexibly choose either class as needed. For example, event-based data can be loaded using the `ContinuousDataset` class by disregarding event timings, and continuous data can be used with the `EpochedDataset` class if it has been preaviously segmented.

Both dataset classes require data in numpy array format, stored in a specified data folder upon instancing. Each subject’s data must be a single array of shape: (trials *×* sensor types *×* sensor locations *×* time samples). More details are available in the online documentation and the prepare data tutorial. Future updates will support the BIDS format ([9, 10]) to streamline data loading and enhance compatibility. Until then, users should use `prepare_data.py` from MEEGNet or follow the prepare data tutorial for proper formatting.

### Model training and testing

The Model class has been designed to be user-friendly and reminiscent of the `Estimator` classes from scikit-learn [11]. It expects data loaded through the dataset classes. The Model class uses PyTorch [12] as its backend, with most of PyTorch’s object methods accessible through the net attribute.

A key design choice in the Model class was the separation of the `feature_extraction` and classification layers into two distinct blocks, accessible via the `feature_extraction` and classification attributes of the `net` object. These objects can be configured separately, which enables the reuse of `feature_extraction` layers across experiments or fine-tuning classification layers for specific tasks.

The `Model` class also supports loading pre-trained models hosted on the Huggingface website or directly from local files. Custom architectures can be created with the `create_net` method, which constructs a network from scratch using specified lists of `feature_extraction` and classification layers.

To provide additional insights into the training process, the library includes a `TrainingTracker` class. This utility automatically gathers key training metrics, such as loss and accuracy, during network training. The collected metrics can then be visualized, offering an intuitive way to monitor model performance and diagnose potential issues.

### Visualization tools

Visualization techniques are essential for understanding neural network decisions, particularly in M/EEG-based neuroscience research. Our package implements two main latent space visualization techniques tailored to the specific needs of ANNs for M/EEG data: Grad-CAM and Guided Backpropagation Saliency. These methods provide neuroscientists with tools to investigate how networks interpret neural signals, facilitating a deeper understanding of brain activity.

Gradient-weighted Class Activation Mapping (Grad-CAM) is particularly effective in highlighting the spatial regions of input data that contribute most significantly to model predictions. Introduced by [13], it generates class-discriminative localization maps by computing the gradient of the target class output *y*^*c*^ (i.e., the score of class *c* before the softmax function) with respect to feature maps from a convolutional layer.

For M/EEG data, Grad-CAM highlights spatial and temporal regions most relevant to the model’s predictions, offering insights into decoding processes. Examples of Grad-CAM are provided in the results section of this paper.

Guided Backpropagation Saliency maps, introduced by [14], focus on input features rather than feature maps, providing a complementary perspective to Grad-CAM. By computing the gradient of the output with respect to the input, saliency maps generate heatmaps that reveal the input features most influential for a prediction. For M/EEG data, this technique highlights temporal and spectral components that drive classification decisions. An example of Guided Backpropagation Saliency applied to ImageNet [15] data is shown in Figure 2, illustrating the approach’s potential for neural data.

**Fig 1.**
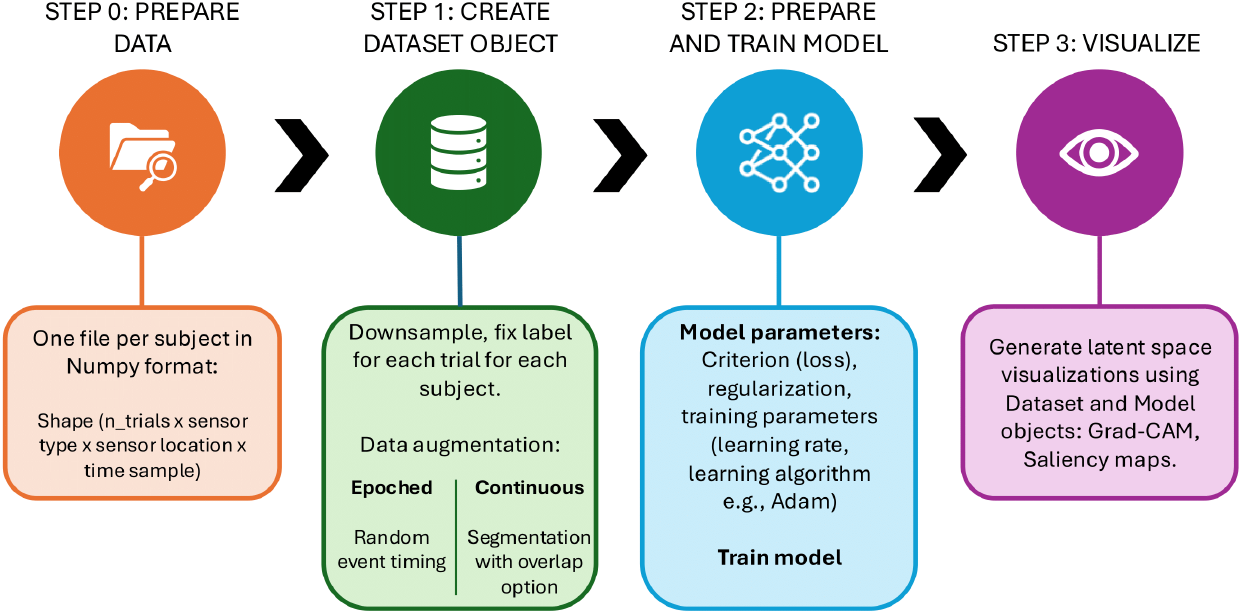
MEEGNet package workflow. Step 0 involves data preparation and problem definition, tasks feasible prior to installing the library. Steps 1–3 are fully supported within MEEGNet. Step 1 entails loading data and creating objects to streamline analysis. Step 2 encompasses deep learning functionalities, including model selection, definition, training, and evaluation. Step 3 (optional) highlights MEEGNet’s strength in visualizing learned representations, enhancing interpretability of classification or regression tasks.

**Fig 2.**
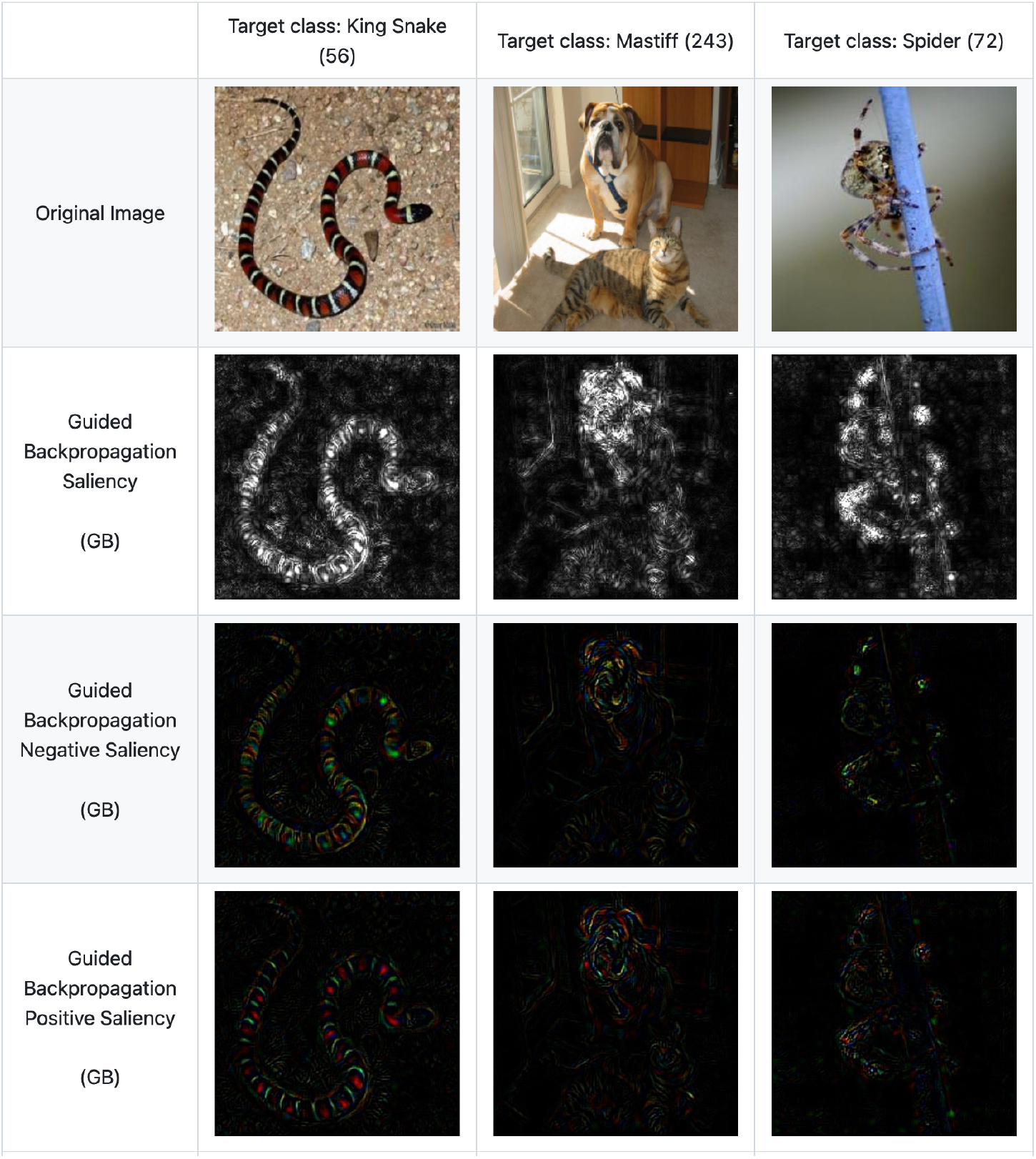
Visualization of saliency maps using ImageNet images, processed by the VGG architectures. The model was pre-trained and adapted for analysis by separating convolutional (features) and fully connected (classifier) layers. Images were normalized with ImageNet mean and standard deviation values. Class labels and corresponding IDs (in parentheses) reflect the ImageNet dataset categorization.

By integrating these visualization methods, our library bridges the gap between black-box ANN architectures and the interpretability required for robust neuroscience research. These tools aim to enhance transparency and foster deeper insights into the neural mechanisms captured by ANNs.

### Tutorials and Documentation

MEEGNet builds upon established, widely-used libraries such as NumPy [16], SciPy [17], PyTorch [12], MNE-Python [18] and Matplotlib [19]. These dependencies are lightweight, well-documented, and prevalent in scientific computing, ensuring ease of installation and compatibility across platforms.

The library is accompanied by comprehensive online documentation hosted on Read the Docs, automatically generated using Sphinx [20] from in-code documentation written in the NumPy style. The documentation provides installation instructions, detailed API references, and step-by-step tutorials, enabling users to quickly familiarize themselves with the library’s features. Additionally, the included tutorials illustrate common use cases, ensuring that both novice and experienced researchers can efficiently integrate MEEGNet into their workflows.

## ANN Architectures

Based on the current literature on ANNs for M/EEG neuroimaging techniques [2], we implemented five architectures: EEGNet [8], VGG-16 [7], MLP [5], AlexNet [6], and our custom-designed MEEGNet. While other ANNs will hopefully be added in the future, we chose to implement these architectures as they span standard and task-specific models, ensuring a versatile performance comparison in MEG decoding tasks. While classical models like MLP, AlexNet, and VGG-16 serve as valuable benchmarks, EEGNet and MEEGNet were developed specifically to address the unique challenges of electrophysiological data, such as spatiotemporal correlations and spectral feature extraction. We will now provide an overview of these five architectures, including their design, application, and relevance to M/EEG analysis. We start with 4 previously available architectures, and then we introduce our custom-designed MEEGNet model.

### MLP

The perceptron, a fundamental building block of artificial neural networks, is a single computational unit that performs linear classification by computing a weighted sum of inputs followed by an activation function. By aggregating multiple perceptrons into layers and connecting them, the multi-layer perceptron (MLP) emerges as a simple yet versatile architecture [21].

In M/EEG research, MLPs are frequently employed as baseline models due to their simplicity and ease of implementation [2]. They provide a straightforward point of comparison for evaluating the performance of more complex architectures. However, MLPs generally lack the interpretability tools and structured latent spaces provided by advanced architectures, which limits, to some extent, their utility for understanding the underlying neural dynamics.

#### AlexNet

AlexNet is a well-established CNN architecture that had a significant impact on deep learning by winning the 2012 ImageNet competition [6]. AlexNet consists of five convolutional layers with relatively large receptive fields (e.g., 11 *×* 11 in the first layer), followed by three FC layers. AlexNet introduced innovations such as Rectified Linear Unit (ReLU) activations, allowing the network to learn non-linear features, and dropout as regularization technique [6].

In the context of MEG research, AlexNet has been adapted as a baseline CNN model, where time and channels are treated analogously to spatial dimensions in images [2, 22–26]. Although it lacks task-specific features for neurophysiological data, as it was primarly designed to perform image classification, its simple yet powerful design makes it a valuable benchmark for exploring spatiotemporal `feature_extraction` in MEG research.

#### VGG-16

VGG, another ImageNet-winning architecture following AlexNet [7], simplifies convolution by stacking 3 *×* 3 receptive fields across deeper layers for hierarchical feature capture. Its popular variant, VGG-16, with 13 convolutional and 3 fully connected layers, offers a strong balance of performance and computational cost compared to deeper VGG models, using fewer parameters.

In MEG research, VGG-16 has been employed as a baseline model, leveraging its depth to capture complex spatiotemporal patterns in neural data. By treating MEG time series and sensor dimensions similarly to spatial dimensions in images, VGG-16 provides a framework for exploring spectral and temporal dynamics [2, 23, 25]. However, its large number of parameters 1 poses challenges in terms of computational cost and overfitting, particularly for smaller MEG datasets.

#### EEGNet

EEGNet is a compact, 3-layer CNN architecture originally designed for EEG-based brain-computer interface (BCI) applications [8, 27]. Widely adopted in EEG research, it serves as a versatile architecture that balances performance and computational efficiency across diverse decoding tasks. EEGNet employs a three-layer convolutional design: depth-wise convolution for spatial filtering, point-wise convolution to reduce dimensionality, and a final 2D convolution for combined spatial and temporal feature extraction.

Drawing inspiration from the DeepConvNet and ShallowConvNet architectures [8], EEGNet is optimized specifically for EEG data, achieving robust results in BCI applications. The architecture has undergone continuous refinement, with EEGNetv4 being the most recent version at the time this paper was written.

To adapt EEGNet for MEG, we introduced modifications that account for the properties of MEG data. Specifically, we added a channel type dimension to handle some MEG device that use three sensors per location (two gradiometers and one magnetometer). For tasks utilizing all sensor types, we tripled the number of learnable parameters to account for the increased complexity.

#### MEEGNet

MEEGNet is a custom-designed CNN architecture tailored for MEG data classification and decoding. Its design was informed by an extensive random search process, optimizing key parameters including depth, filter size, and number of channels, to achieve a balance between decoding accuracy and interpretability. Note that the hyperparameter optimization process was performed using MEG resting-state data (as detailed in the MEG data sets section of this paper) on a biological sex classification task. This task was not included in the benchmarking data sets in order to avoid risks of double dipping.

As detailed in Table 2, MEEGNet begins with a spatial convolution layer that aggregates information across MEG sensors at each time point. This is followed by temporal convolution layers, which capture spectral and temporal dynamics at multiple scales. Max pooling is applied at specific stages to reduce the temporal dimension, maintaining computational efficiency while preserving critical features. The architecture concludes with two fully connected (FC) layers, which map the extracted features to class probabilities, making it adaptable for both binary and multi-class classification tasks.

**Table 1.** Number of parameters for each network included in the MEEGNet Library.

| Network | Parameters |
| --- | --- |
| MEEGNet (ours) | 32.6M |
| EEGNet | 14.3k |
| VGG | 339M |
| MLP | 12.2M |

**Table 2.** MEEGNet Architecture: Description of all layers.

| Name | Layer Type | Description | Output Dimensions |
| --- | --- | --- | --- |
| **Feature Extraction Layers** |  |  |  |
| Conv1 | Conv2d | Input channels: 3, Output: 100,<br>Kernel: (102, 1) | $100 \times 1 \times 400$ |
| | ReLU | Activation function | $100 \times 1 \times 400$ |
| Conv2 | Conv2d | Input channels: 100, Output: 200,<br>Kernel: (1, 9) | $200 \times 1 \times 392$ |
| | MaxPool2d | Kernel: (1, 20) | $200 \times 1 \times 373$ |
| | ReLU | Activation function | $200 \times 1 \times 373$ |
| Conv3 | Conv2d | Input channels: 200, Output: 200,<br>Kernel: (1, 9) | $200 \times 1 \times 365$ |
| | MaxPool2d | Kernel: (1, 20) | $200 \times 1 \times 346$ |
| | ReLU | Activation function | $200 \times 1 \times 346$ |
| Conv4 | Conv2d | Input channels: 200, Output: 100,<br>Kernel: (1, 9) | $100 \times 1 \times 338$ |
| | MaxPool2d | Kernel: (1, 20) | $100 \times 1 \times 319$ |
| | ReLU | Activation function | $100 \times 1 \times 319$ |
|  | Flatten | Flatten spatial dimensions to vector | 31900 |
|  | Dropout | Dropout probability: 0.5 | 31900 |
| **Classification Layers** |  |  |  |
| FC4 | Linear | Input features: 31900,<br>Output: 1000 | 1000 |
| FC6 | Linear | Input features: 1000,<br>Output: 2 | 2 |

A visual representation of the MEEGNet architecture is provided in Figure 3, illustrating the data flow through the `feature_extraction` and classification blocks. The modularity of MEEGNet is a key strength, allowing researchers to adapt the architecture to specific datasets or tasks. For instance, the `feature_extraction` layers can be reused across experiments, while the classification layers can be fine-tuned for task-specific requirements.

**Fig 3.**
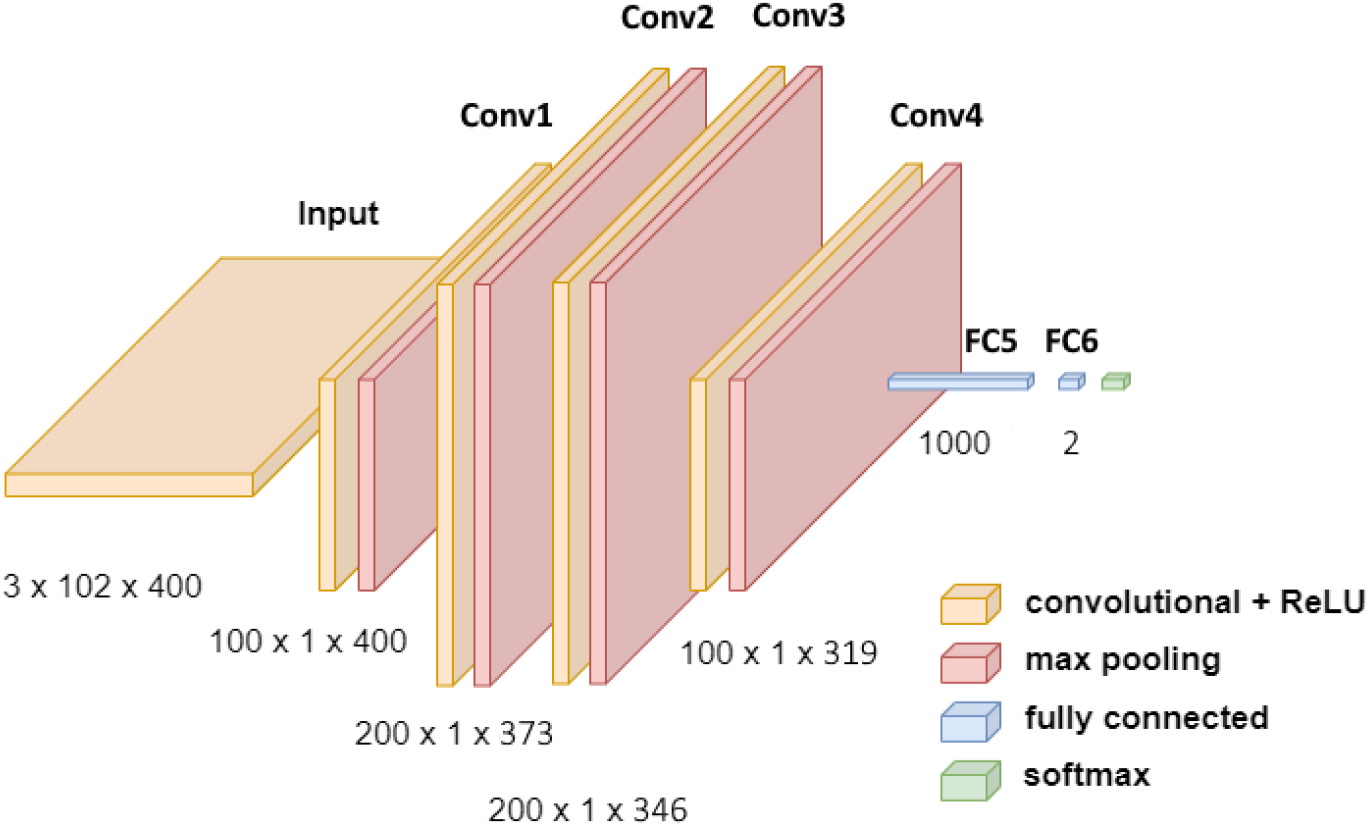
Design of our MEEGNet architecture. This MEEGNet example has four convolutional layers and two outputs for binary classification. This network is used for instance in the auditory vs visual stimulus classification task. See Table 2 for details of the MEEGNet architecture.

### Validation and benchmarking

#### MEG datasets

We used open-access MEG data from the Cambridge Centre for Ageing and Neuroscience (Cam-CAN) [28], consisting of resting-state recordings and a passive audio-visual perception task. This dataset was selected for its large sample size (n=643) and well-documented preprocessing pipelines, ideal for testing our neural network implementations.

Resting-state data included 520-second MEG recordings per subject, downsampled from raw data to 200 Hz and epoched into 2-second trials, yielding samples of shape 3 × 102 × 400 (sensor types: gradiometers and magnetometers; MEG channels; time samples). After removing bad trials, the final dataset comprises 120,000 trials across 643 subjects. For the passive task, 2-minute sessions presented visual checkerboards or auditory tones every second in random order, with data resampled from 1000 Hz to 500 Hz and epoched into 800 ms trials, also shaped 3 × 102 × 400. Post-preprocessing, this stimulus classification dataset totals 677,730 trials. When combining resting-state and passive task data for tasks like stimulus vs. resting-state classification, we resampled the resting-state recordings from raw data to 500 Hz to match the passive task sampling rate.

#### Classification performance evaluation

We evaluated the performance of our library using four classification tasks: (1) Auditory vs. visual stimulus classification, (2) Auditory stimulus vs. resting state, (3) Visual stimulus vs. resting state, and (4) Age classification (20–29 to 80–89) from resting-state data. Task (1) serves as a benchmark for validating latent space visualization techniques, leveraging the well-established spatial, temporal, and spectral dynamics of auditory and visual stimuli [29, 30]. Tasks (2) and (3) test the network’s ability to distinguish stimulus-evoked activity from resting-state signals, which exhibit distinct neural dynamics. Task (4) assesses the network’s capacity to classify subjects into decade-based age categories, posing a multi-class challenge and testing robustness to inter-subject variability.

Performance metrics included validation and test accuracy, computation time, and the number of epochs to convergence, using an 80 *−* 10 *−* 10 train/validation/test split for consistency across networks. Models were initialized with default parameters or those from their original implementations and trained using the Adam optimizer [2] with cross-entropy loss. Early stopping was applied after 10 epochs without validation loss improvement to prevent overfitting, and a fixed random state ensured reproducibility.

To benchmark our deep learning models, we also evaluated a Logistic Regression baseline. Since classical machine learning struggles with raw data’s high dimensionality, we computed Power Spectral Density (PSD) values using the Welch method [31], averaged over frequency bands: delta (0.5–4 Hz), theta (4–8 Hz), alpha (8–12 Hz), low beta (12–30 Hz), high beta (30–60 Hz), low gamma (60–90 Hz), and high gamma (90–120 Hz). This produced a 3 *×* 102 *×* 7 feature matrix per trial. The model was optimized via a 100-iteration random search (see Table 3) with nested leave-50-subjects-out cross-validation, selecting the best configuration of sensor type and location based on validation accuracy.

**Table 3.** Hyperparameters tested for Logistic Regression in the random search. Each hyperparameter was varied across the specified values during optimization to identify the best configuration for each MEG classification task. C is the inverse of regularization strength, controlling the trade-off between fitting the training data and preventing overfitting (smaller values increase regularization). Penalty specifies the regularization type (L1 or L2), influencing sparsity or coefficient shrinkage during training. Max Iterations defines the maximum number of iterations for the solver to converge, serving as a stopping criterion when training. Tolerance sets the threshold for convergence, determining when the optimization process stops based on the change in the loss function. Fit Intercept indicates whether to include an intercept term in the model, allowing the regression line to shift away from the origin if set to True. Class Weight adjusts the influence of each class, with “balanced” compensating for class imbalance by weighting inversely to class frequency, or “None” applying equal weights.

| Hyperparameter | Values Tested |
| --- | --- |
| C | {0.01, 0.046, 0.215, 1.0, 4.64, 21.54, 100.0} (logarithmically spaced) |
| Penalty | {l1, l2} |
| Max Iterations | {100, 200, 500, 1000} |
| Tolerance (tol) | {$10^{-4}$, $10^{-3}$, $10^{-2}$, $10^{-1}$} |
| Fit Intercept | {True, False} |
| Class Weight | {None, balanced} |

## Results

In this section, we present findings on task performance and latent space visualizations as detailed in the Design and Implementation section.

### Network Performances

The results of our experiments, summarized in Table 4, demonstrate the effectiveness of MEEGNet in classifying MEG signals across various tasks, showcasing its robust performance, computational efficiency, and balanced parameter usage.

**Table 4.** Comparison of classification performance between MEEGNet (ours), other commonly used architectures in the M/EEG literature (EEGNet, VGG, and MLP), and a baseline Logistic Regression (LR) model. The table reports test accuracy (T Acc), test loss (T Loss), validation accuracy (V Acc), validation loss (V Loss), number of epochs (Epoch), and training time (Time) for each architecture across four classification tasks: Visual Stimulus vs. Resting State, Auditory Stimulus vs. Resting State, and age classification (7-class problem).

| Task | Network | T Acc | T Loss | V Acc | V Loss | Epoch | Time (hh:mm) |
| --- | --- | --- | --- | --- | --- | --- | --- |
| **Auditory vs. Visual** | MEEGNet (ours) | **92.70** | **0.1776** | **94.24** | **0.1391** | 149 | **02:42** |
|  | EEGNet | 91.14 | 0.2111 | 92.47 | 0.1804 | 256 | 06:00+ |
|  | VGG | 91.36 | 0.2275 | 93.05 | 0.1702 | 94 | 03:51 |
|  | MLP | 90.79 | 0.2232 | 92.39 | 0.1836 | 303 | 03:03 |
|  | Baseline (LR) | 63.56 | n/a | 63.93 | n/a | n/a | 00:17 |
| **Visual vs. Rest** | MEEGNet (ours) | 93.06 | 0.3822 | 94.92 | 0.1373 | 47 | **00:37** |
|  | EEGNet | 94.91 | 0.1508 | 96.47 | 0.0998 | 178 | 02:40 |
|  | VGG | **98.30** | **0.0593** | **98.82** | **0.0366** | 58 | 02:39 |
|  | MLP | 60.36 | 0.6411 | 63.44 | 0.6144 | 191 | 02:05 |
|  | Baseline (LR) | 74.21 | n/a | 72.38 | n/a | n/a | 00:09 |
| **Auditory vs. Rest** | MEEGNet (ours) | 94.27 | 0.3502 | 94.68 | 0.1428 | 58 | **00:43** |
|  | EEGNet | 95.62 | 0.1249 | 97.12 | 0.0801 | 186 | 03:31 |
|  | VGG | **98.45** | **0.0747** | **98.76** | **0.0350** | 68 | 03:06 |
|  | MLP | 50.00 | 0.6931 | 50.00 | 0.6931 | 214 | 01:59 |
|  | Baseline (LR) | 64.37 | n/a | 64.50 | n/a | n/a | 00:12 |
| **Age (7-class)** | MEEGNet (ours) | **23.60** | 1.8360 | **22.70** | **1.8878** | 6 | **0:33** |
|  | EEGNet | 21.06 | **1.7361** | 19.60 | 1.9535 | 10 | 02:08 |
|  | VGG | 22.63 | 1.8194 | 21.70 | 1.9429 | 7 | 00:58 |
|  | MLP | 9.29 | 1.9681 | 22.11 | 1.9346 | 20 | 01:53 |
|  | Baseline (LR) | 15.22 | n/a | 16.36 | n/a | n/a | 01:22 |

In first classification task, Auditory vs. Visual Stimulus, MEEGNet outperforms all other architectures, achieving a test accuracy of 92.70% and a test loss of 0.1776. Notably, it converges in 149 epochs, compared to EEGNet’s 256, and trains more than twice as fast (02:42 vs. 06:00+). The baseline model achieves a test accuracy of 63.56%, again underscoring the importance of neural networks’ ability to learn task-relevant representations directly from the data.

For the second task, Visual Stimulus vs. Resting State, MEEGNet achieves a test accuracy of 93.06% with a test loss of 0.3822. While VGG outperforms MEEGNet in accuracy (98.30% vs. 93.06%) and loss (0.0593 vs. 0.3822), MEEGNet trains significantly faster (00:37 vs. 02:39) and converges in fewer epochs (47 vs. 58). The baseline Logistic Regression model achieves a test accuracy of 74.21%, which highlights the added value of neural networks’ representation learning capabilities. While LR relies on manually extracted features, neural networks learn representations directly from the raw data, allowing them to better capture the spatiotemporal dynamics of MEG signals.

In the Auditory Stimulus vs. Resting State classification task, MEEGNet again demonstrates competitive performance, achieving a test accuracy of 94.27% and a test loss of 0.3502. Although VGG attains higher accuracy (98.45%) and lower loss (0.0747), MEEGNet completes training in less than a third of the time (00:43 vs. 03:06) while maintaining a significantly smaller parameter count (32.6M vs. 339M). The baseline model achieves a test accuracy of 64.37%, reflecting the limited scope of manually extracted features compared to the rich feature representations learned by neural networks.

For the exploratory Age Classification task, MEEGNet achieves the best overall decoding performance, with a test accuracy of 23.60% and a validation accuracy of 22.70%. As this is a 7-class classification problem with slight class unbalance, the corrected theoretical chance level is 17.98%. Although performance on this task is inherently challenging due to the fine-grained nature of age group predictions, MEEGNet requires only six epochs to converge and completes training in 00:33, compared to VGG’s 00:58 and EEGNet’s 02:08. The baseline model achieves a test accuracy of 15.22%, further emphasizing the challenges of age prediction and the limited adaptability of LR with predefined features.

Across all tasks, MEEGNet showcases its strengths in decoding MEG data with competitive accuracy and fast training times. The relatively modest performance of the Logistic Regression baseline underscores the critical role of representation learning in attaining superior performance outcomes, particularly for complex spatiotemporal data like MEG. These results position MEEGNet as a robust and reliable architecture for MEG-based classification tasks, balancing performance with practical usability for researchers working in the field.

### Visualization techniques

Visualization techniques were assessed using EEGNet on the stimulus classification task, an ideal benchmark due to its simplicity and well-characterized auditory-visual responses. Figure 4 shows event-related potentials (ERPs) averaged across subjects and trials as a reference. EEGNet was selected for its lightweight design and Grad-CAM compatibility, unlike VGG’s large size, which slows visualization, or MEEGNet’s unique architecture, which does not optimally highlight these techniques.

**Fig 4.**
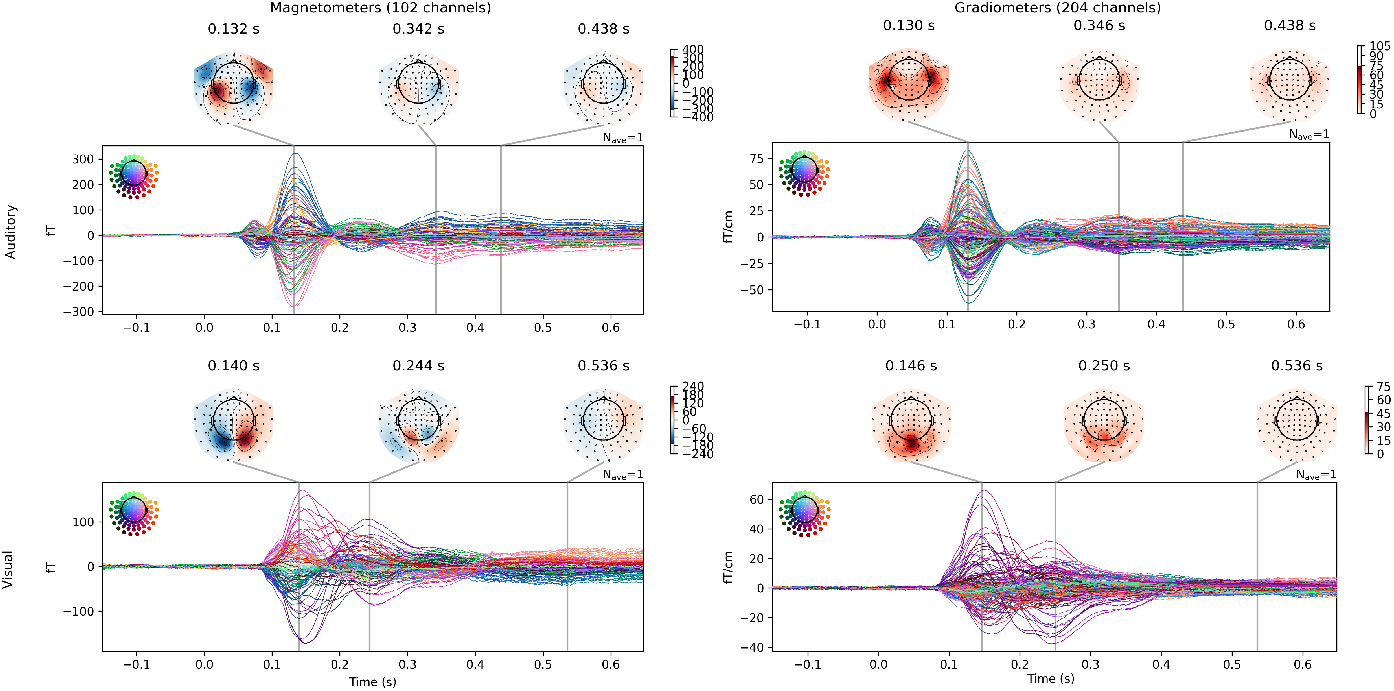
Event-Related Potentials (ERPs) for auditory (first row) and visual (second row) stimuli during the audio-visual passive task from the Cam-CAN dataset. ERPs are averaged across all trials and subjects, with stimulus onset at t=0s. Trials were baseline-corrected using the pre-stimulus interval. Topomaps above each ERP provide a spatial representation of stimulus-evoked neural activity at three automatically detected peaks, highlighting the spatial distribution of the ERP across sensor locations.

#### Grad-CAM Visualization

Gradient-weighted Class Activation Mapping (Grad-CAM) visualizes key input data regions influencing neural network predictions by generating class-discriminative heatmaps from gradients of a class score relative to convolutional feature maps. For MEG data, it reveals spatial and temporal features critical to classification, linking predictions to neurophysiological processes.

Figure 5 illustrates Grad-CAM activations generated by the EEGNet architecture for the Visual Stimulus vs. Auditory Stimulus classification task. Left-column heatmaps, averaged across trials and subjects per stimulus class, show distinct neural activity patterns for visual and auditory stimuli. Right-column topographic maps, derived from the first PCA component of Grad-CAM at peak activation, pinpoint sensor locations crucial for accurate stimulus classification. For the auditory stimulus, maximum activation occurs at *t* = 0.155*s*, and for the visual stimulus at *t* = 0.121*s*. These timings align with known neural response dynamics, as ERPs typically peak at *t* = 0.131*s* for auditory (Fig. 4), and at *t* = 0.143*s* for visual stimuli. The topomaps highlight strong activations in occipital sensors for visual stimuli and temporal sensors for auditory stimuli, consistent with the known roles of the visual [32] and auditory cortices [33].

**Fig 5.**
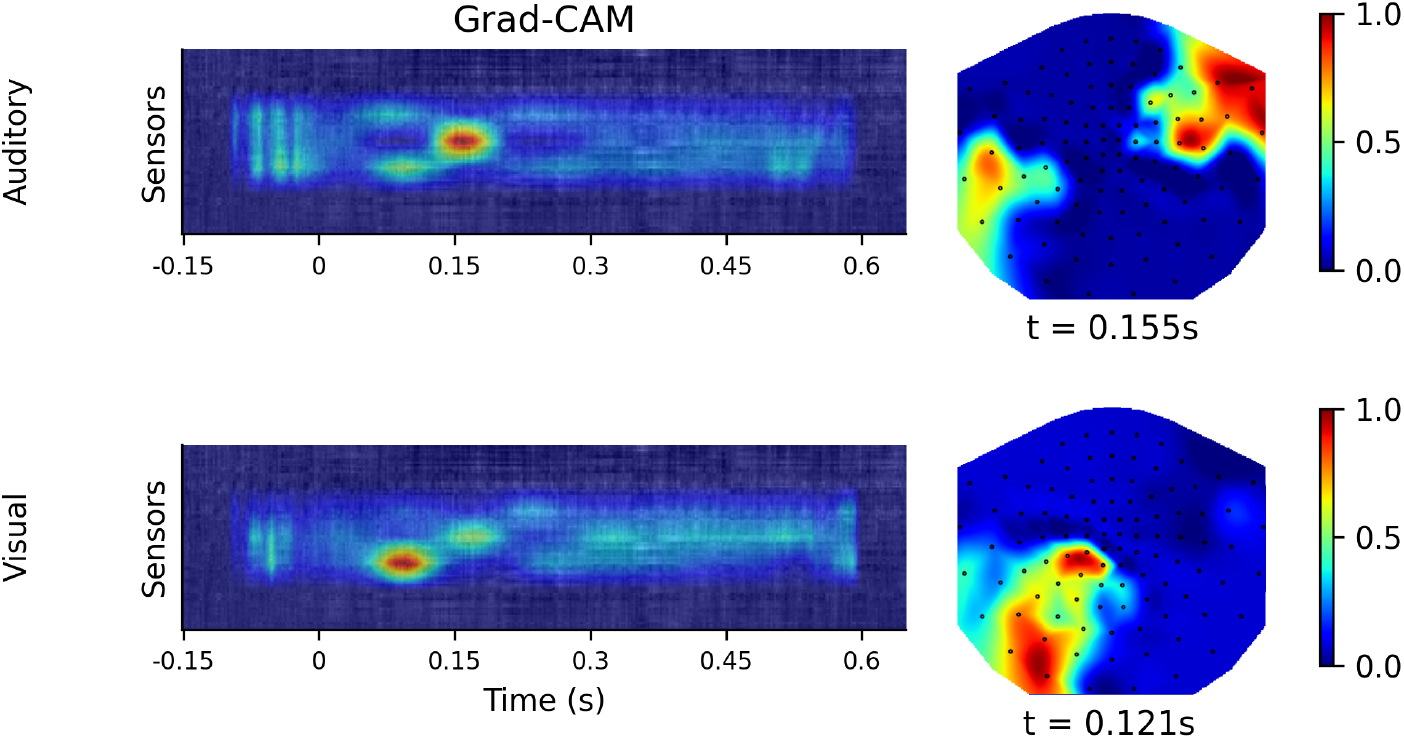
Grad-CAM activation maps for the final layer of the EEGNet architecture during the Visual vs. Auditory Stimulus classification task. The left column shows example trials with Grad-CAM activations averaged across all trials and subjects, limited to predictions with confidence *≥* 90%. The right column presents topographic maps derived from the first principal component of the Grad-CAM masks across all subjects (representing 55.57% and 71.85% of explained variance for auditory, first row, and visual, second row, stimulus, respectively), captured at the time point of maximum activation. The first row corresponds to auditory stimulus trials, while the second row corresponds to visual stimulus trials.

Grad-CAM visualizations, combined with PCA-derived topomaps, enhance understanding of neural network activations. This pairing highlights key temporal and spatial features for predictions while aligning them with established neuroanatomical and functional principles.

#### Saliency maps

Saliency maps visualize input features influencing neural network predictions by projecting output gradients onto heatmaps. For MEG data, they reveal key temporal and spatial features driving classification. They can be computed independently for each sensor type, allowing researchers to evaluate magnetometer and gradiometer contributions to network predictions. This reveals how different sensor modalities aid classification.

Figure 6 illustrates saliency maps generated by the EEGNet architecture for the Auditory vs. Visual Stimulus classification task. The heatmaps averaged across all trials and subjects for each stimulus class, provide a robust estimation of the time points and sensor locations most influential in the network’s predictions. As anticipated, channels associated with auditory processing exhibit higher saliency for auditory stimuli, while visual-related channels dominate for visual stimuli. This pattern aligns with established knowledge about auditory and visual processing pathways in the brain ([32, 33]).

**Fig 6.**
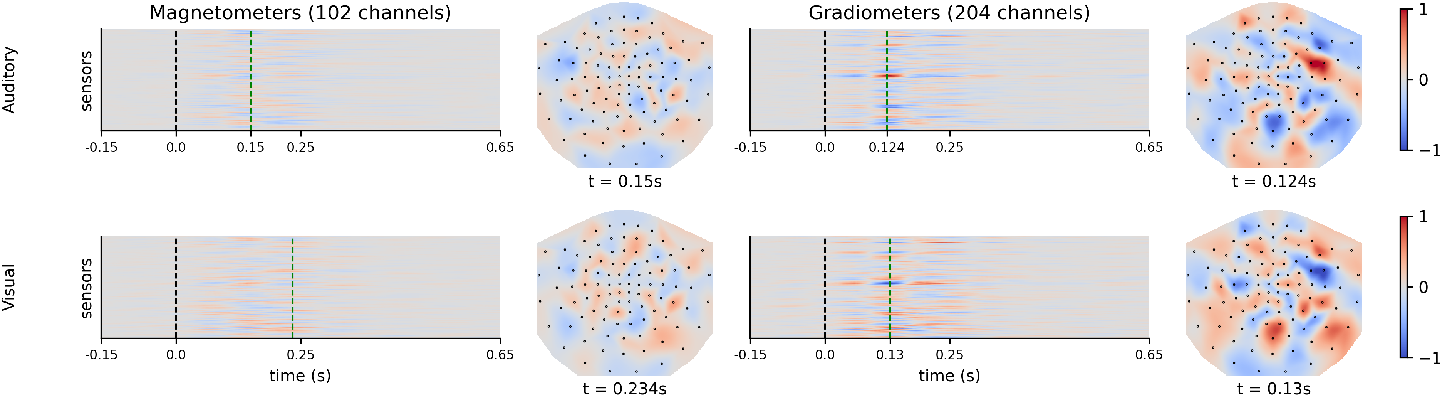
Guided backpropagation saliency maps for the EEGNet architecture during the stimulus classification task. Saliency maps and topographic representations for auditory and visual stimuli predictions with *≥* 90% confidence. Magnetometer (left) and gradiometer (right) saliency maps, with the x-axis representing time (s) and the y-axis representing sensor locations. Green dashed lines indicate peak saliency timing; black dashed lines mark stimulus onset (t=0s). Topomaps display the spatial distribution of peak saliency at key time points.

The topographic representations in Figure 6 further enhance interpretability by summarizing the spatial distribution of peak saliency across the sensor array. For auditory stimuli, the peak saliency timing for magnetometers is observed at *t* = 0.15*s*, aligning with the first auditory ERP peak. For gradiometers, the peak timing occurs slightly earlier at *t* = 0.124*s*. For visual stimuli, magnetometer saliency peaks at *t* = 0.234*s*, corresponding approximately to the second visual ERP peak at *t* = 0.247*s* (See Fig. 4), while gradiometer saliency peaks earlier at *t* = 0.13*s*. These results demonstrate the network’s ability to capture neurophysiologically relevant features with high precision and modality-specific detail.

Saliency maps link computational predictions to neurophysiology. MEEGNet offers built-in tools to generate these visualizations across architectures, helping researchers analyze sensor contributions and network activations in relation to brain dynamics.

## Availability and Future Directions

The MEEGNet library is available as a free, open-source package under the MIT license on GitHub and PyPI, installable via pip. It includes online documentation with installation guides, tutorials, and visualization examples. We encourage feedback to enhance flexibility and support diverse use cases. Future enhancements for MEEGNet include:

- **Expanding Model Architectures:** Integrate additional architectures, such as LF-CNN and VAR-CNN [34], and advanced foundation models for M/EEG data, like the foundation GPT Model for MEG [35], EEG [36], and Nested Deep Learning Models for brain signal data [37]. This will ensure MEEGNet remains at the forefront of the field.
- **Improving our MEEGNet Architectures:** As more research using M/EEG-tailored ANN architectures emerges, it is crucial to incorporate new building blocks and insights to keep our architecture competitive. A MEEGNetv2 architecture is already in development and will soon be added to the package, replacing the original one, which will be retained as a legacy option.
- **Data Augmentation Techniques:** Implement robust data augmentation methods to mitigate data scarcity and enhance model generalization.
- **Advanced Visualization Tools:** Expand visualization techniques (e.g., Filter Visualizations) to enhance interpretability, leveraging tools from other domains.
- **Integration into Larger Projects:** Consider merging MEEGNet into larger platforms to maximize impact and usability, as well as distribute maintenance work.
- **BIDS Compatibility:** Implement compatibility with the Brain Imaging Data Structure (BIDS) format to streamline data loading and preprocessing.
- **Enhanced Tutorials and Documentation:** Generate more examples and tutorials, refining resources based on user feedback.

MEEGNet’s open-source nature aligns with the broader movement toward open science in neuroscience [38]. By making the library freely available, we aim to foster collaboration, encourage reproducibility, and contribute to the democratization of advanced analytical tools, ensuring MEEGNet is a valuable resource for the neuroscience community.

## Conflict of interest

The authors declare that the research was conducted in the absence of any commercial or financial relationships that could be construed as a potential conflict of interest.

## A. Supplementary Materials

### A.1 Supplementary Insights into MEEGNet’s Development and Performance

The development of MEEGNet represents a significant step toward providing neuroscientists with a suite of intuitive, easy-to-use tools for ANN-driven electrophysiological data analysis, with a specific emphasis on decoding and interpretability. The primary goal of this project is to bridge the gap between advanced artificial neural networks (ANNs) and their practical application in neuroscience by offering a library that is accessible, modular, and adaptable to the specific needs of electrophysiological data.

MEEGNet was designed to streamline ANN-powered MEG data analysis through an open-source efficient pipeline. The library includes functionalities for data preprocessing, model training, and evaluation. Most importantly, it integrates visualization tools that allow users to explore the latent spaces of their models, providing insight into how decisions are made and which features are critical for decoding. These capabilities address longstanding challenges of interpretability in ANN-based approaches, especially for MEG and EEG data.

The validation experiments conducted in this study demonstrate that MEEGNet reliably decodes electrophysiological data while offering insights through its visualization tools. The results show that the library achieves strong performance across various tasks and provides interpretable outputs that highlight meaningful patterns in the data. These findings underscore the utility of MEEGNet as a robust and reliable tool for neuroscience research.

A key strength of MEEGNet lies in its modular design, which facilitates customization and contribution. Users can easily integrate new components and models, making the library a flexible platform for innovation. This modularity ensures that MEEGNet can evolve alongside advancements in the field, accommodating new techniques and addressing emerging challenges.

However, there are limitations to this approach. The effectiveness of MEEGNet depends on the quality and quantity of the data it is applied to, and its performance may be constrained by challenges common to neural networks, such as data scarcity and overfitting. Additionally, while the library emphasizes interpretability, some aspects of ANN decision-making remain inherently opaque. Further development of visualization and explainability techniques is needed to fully address these limitations.

### A.2 MEEGNet Hyperparameter search

**Table 5.** Hyperparameters tested for the creation of the MEEGNet Network. Random Search was used to select random set of these parameters to test on a sex identification classification problem using the CamCAN dataset as described in the methods section. The Number of channels parameter was used as size of the smallest layers, other layers were scaled from this number by a factor of two. For example, in the final architecture, depth of 4 was selected with number of channels of 100, leading to 4 layers: one of 100 channels followed by two of 200 channels and another 100-channel layer.

| Hyperparameter | Values Tested |
| --- | --- |
| Depth | {3, 4, 5, 6, 7, 8, 9} |
| Filter size | {3, 5, 7, 9, 12, 15} |
| Number of Channels | {25, 50, 100, 200, 400, 800} |
| Number of Linear Units | {100, 250, 500, 1000, 2000, 4000} |
| Mini Batch Size | {16, 32, 64, 128, 256} |

## Notes

### Competing Interest Statement

The authors have declared no competing interest.

## References

1. LeCun Y, Bengio Y, Hinton G. Deep learning. nature. 2015;521(7553):436–444.

2. Dehgan A, Rish I, Jerbi K. Artificial Neural Networks for Magnetoencephalography: A review of an emerging field. Public Library of Science San Francisco, CA USA; 2025.

3. Zubarev I, Vranou G, Parkkonen L. MNEflow: Neural networks for EEG/MEG decoding and interpretation. SoftwareX. 2022;17:100951.

4. Schirrmeister RT, Springenberg JT, Fiederer LDJ, Glasstetter M, Eggensperger K, Tangermann M, et al. Deep learning with convolutional neural networks for EEG decoding and visualization. Human Brain Mapping. 2017;doi:10.1002/hbm.23730.

5. Almeida LB. Multilayer perceptrons. In: Handbook of Neural Computation. CRC Press; 2020. p. C1–2.

6. Krizhevsky A, Sutskever I, Hinton GE. Imagenet classification with deep convolutional neural networks. Advances in neural information processing systems. 2012;25.

7. Simonyan K, Zisserman A. Very deep convolutional networks for large-scale image recognition. arXiv preprint arXiv:14091556. 2014;.

8. Lawhern VJ, Solon AJ, Waytowich NR, Gordon SM, Hung CP, Lance BJ. EEGNet: a compact convolutional neural network for EEG-based brain–computer interfaces. Journal of neural engineering. 2018;15(5):056013.

9. Niso G, Gorgolewski KJ, Bock E, Brooks TL, Flandin G, Gramfort A, et al. MEG-BIDS, the brain imaging data structure extended to magnetoencephalography. Scientific data. 2018;5(1):1–5.

10. Pernet CR, Appelhoff S, Gorgolewski KJ, Flandin G, Phillips C, Delorme A, et al. EEG-BIDS, an extension to the brain imaging data structure for electroencephalography. Scientific data. 2019;6(1):103.

11. Pedregosa F, Varoquaux G, Gramfort A, Michel V, Thirion B, Grisel O, et al. Scikit-learn: Machine Learning in Python. Journal of Machine Learning Research. 2011;12:2825–2830.

12. Paszke A, Gross S, Massa F, Lerer A, Bradbury J, Chanan G, et al. PyTorch: An Imperative Style, High-Performance Deep Learning Library. In: Advances in Neural Information Processing Systems 32. Curran Associates, Inc.; 2019. p. 8024–8035. Available from: http://papers.neurips.cc/paper/9015-pytorch-an-imperative-style-high-performance-deep-learning-library.pdf.

13. Selvaraju RR, Cogswell M, Das A, Vedantam R, Parikh D, Batra D. Grad-CAM: visual explanations from deep networks via gradient-based localization. International journal of computer vision. 2020;128:336–359.

14. Simonyan K. Deep inside convolutional networks: Visualising image classification models and saliency maps. arXiv preprint arXiv:13126034. 2013;.

15. Deng J, Dong W, Socher R, Li LJ, Li K, Fei-Fei L. Imagenet: A large-scale hierarchical image database. In: 2009 IEEE conference on computer vision and pattern recognition. Ieee; 2009. p. 248–255.

16. Harris CR, Millman KJ, van der Walt SJ, Gommers R, Virtanen P, Cournapeau D, et al. Array programming with NumPy. Nature. 2020;585(7825):357–362. doi:10.1038/s41586-020-2649-2.

17. Virtanen P, Gommers R, Oliphant TE, Haberland M, Reddy T, Cournapeau D, et al. SciPy 1.0: Fundamental Algorithms for Scientific Computing in Python. Nature Methods. 2020;17:261–272. doi:10.1038/s41592-019-0686-2.

18. Gramfort A, Luessi M, Larson E, Engemann DA, Strohmeier D, Brodbeck C, et al. MEG and EEG Data Analysis with MNE-Python. Frontiers in Neuroscience. 2013;7(267):1–13. doi:10.3389/fnins.2013.00267.

19. Hunter JD. Matplotlib: A 2D graphics environment. Computing in Science & Engineering. 2007;9(3):90–95. doi:10.1109/MCSE.2007.55.

20. Brandl G. Sphinx documentation. URL http://sphinx-docorg/sphinxpdf. 2021;.

21. Goodfellow I, Bengio Y, Courville A, Bengio Y. Deep learning. vol. 1. MIT press Cambridge; 2016.

22. Dima DC, Perry G, Singh KD. Spatial frequency supports the emergence of categorical representations in visual cortex during natural scene perception. NeuroImage. 2018;179:102–116.

23. Giari G, Leonardelli E, Tao Y, Machado M, Fairhall SL. Spatiotemporal properties of the neural representation of conceptual content for words and pictures–an MEG study. Neuroimage. 2020;219:116913.

24. Rajaei K, Mohsenzadeh Y, Ebrahimpour R, Khaligh-Razavi SM. Beyond core object recognition: Recurrent processes account for object recognition under occlusion. PLoS computational biology. 2019;15(5):e1007001.

25. Giovannetti A, Susi G, Casti P, Mencattini A, Pusil S, López ME, et al. Deep-MEG: spatiotemporal CNN features and multiband ensemble classification for predicting the early signs of Alzheimer’s disease with magnetoencephalography. Neural Computing and Applications. 2021;33(21):14651–14667.

26. Dash D, Ferrari P, Wang J. Decoding imagined and spoken phrases from non-invasive neural (MEG) signals. Frontiers in neuroscience. 2020;14:290.

27. Waytowich N, Lawhern VJ, Garcia JO, Cummings J, Faller J, Sajda P, et al. Compact convolutional neural networks for classification of asynchronous steady-state visual evoked potentials. Journal of neural engineering. 2018;15(6):066031.

28. Shafto MA, Tyler LK, Dixon M, Taylor JR, Rowe JB, Cusack R, et al. The Cambridge Centre for Ageing and Neuroscience (Cam-CAN) study protocol: a cross-sectional, lifespan, multidisciplinary examination of healthy cognitive ageing. BMC neurology. 2014;14(1):1–25.

29. Creel DJ. Visually evoked potentials. Handbook of clinical neurology. 2019;160:501–522.

30. Picton TW, Hillyard SA, Krausz HI, Galambos R. Human auditory evoked potentials. I: Evaluation of components. Electroencephalography and clinical neurophysiology. 1974;36:179–190.

31. Solomon Jr OM. PSD computations using Welch’s method. NASA STI/Recon Technical Report N. 1991;92:23584.

32. Wandell BA, Dumoulin SO, Brewer AA. Visual field maps in human cortex. Neuron. 2007;56(2):366–383.

33. Zatorre RJ, Bouffard M, Ahad P, Belin P. Where is’ where’in the human auditory cortex? Nature neuroscience. 2002;5(9):905–909.

34. Zubarev I, Zetter R, Halme HL, Parkkonen L. Adaptive neural network classifier for decoding MEG signals. NeuroImage. 2019;197. doi:10.1016/j.neuroimage.2019.04.068.

35. Csaky R, van Es MW, Jones OP, Woolrich M. Foundational GPT Model for MEG. arXiv preprint arXiv:240409256. 2024;.

36. Cui W, Jeong W, Thölke P, Medani T, Jerbi K, Joshi AA, et al. Neuro-gpt: Towards a foundation model for eeg. In: 2024 IEEE International Symposium on Biomedical Imaging (ISBI). IEEE; 2024. p. 1–5.

37. Wei F, Mo J, Zhang K, Shen H, Nagarajan S, Jiang F. Nested Deep Learning Model Towards A Foundation Model for Brain Signal Data. arXiv preprint 241003191. 2024;.

38. Paret C, Unverhau N, Feingold F, Poldrack RA, Stirner M, Schmahl C, et al. Survey on open science practices in functional neuroimaging. NeuroImage. 2022;257:119306.

